# Reading Outside the Lines: A Systematic Approach for Detecting Bias in Scientific Communications

**DOI:** 10.1101/2024.06.12.598649

**Authors:** Melisa Osborne, TJ McKenna, Felicity Crawford, Theresa Rueger, Barkha Shah, Mae Rose Gott, Adam Labadorf

**Affiliations:** Graduate Program in Bioinformatics, Boston University, Boston, MA; Wheelock College of Education and Human Development, Boston University, Boston, MA; School of Natural and Environmental Sciences, Newcastle University, UK; CAS Department of Biology, Boston University, Boston, MA; Graduate Program in Bioinformatics & Department of Neurology, Boston University Chobanian & Avedisian School of Medicine, Boston, MA

## Abstract

Consciousness of the social impact of science and the potential biases of its authors is critical to understanding, interpreting, and using scientific findings responsibly. This is especially true for sciences concerned with human health and behavior, where societal and unconscious biases may reinforce existing inequities and discriminatory practices. Considering backgrounds and biases, we may notice bias influencing scientists’ methodological choices and conclusions, even when a work is otherwise scientifically sound. To this end, we created the pedagogical tool Finding inEquity in LIterature and eXperimentation (FELIX). FELIX is a tool that systematizes the detection of bias and subjectivity in scientific communications by using a three-phase progression of (i) Annotation, (ii) Analysis and (iii) Synthesis, where students form a unified argument about the text with a focus on its relationship to social or ethical context. Results from a mixed methods approach indicated the efficacy of our approach in supporting student learning related to reading comprehension, critical thinking skills, and in understanding the social and ethical implications of the research they were reading. We put forward FELIX as a universal method for training students in the reading of scientific communications and as a tool for addressing systemic inequities in science and science education.

## INTRODUCTION

Since the establishment of the field of biology, scientific work has both been impacted by the society in which it is practiced and has in turn had an impact on policy and society. The example of taxonomic hierarchies of human groups ideated in the late 1700’s by Linnaeus, Blumenbach, and others was a reflection of societal views on race and in turn a means for continued justification of colonialism and slavery around the globe (Saini, 2019; Muller-Wille, 2014). The science-and-society feedback loop regarding race thus having been established - assumptions about biological differences between people from different so-called racial groups have been maintained throughout the history of science and have been hard to remove, even in the light of their pseudoscientific nature. The history of biology and genetics is littered with examples of how devaluation of groups lower in the social hierarchy has led to unethical scientific practices and personal harm. This is most apparent in considering harms generated during the Eugenic era in America and Europe in the 20^th^ century, including the Tuskegee incident (1930’s -1970s) and cruel scientific experimentation carried out by Nazi Germany (Washington, 2008; Gould, 1996; Okrent, 2020). Changes to scientific ethical standards post World War II directly addressed the horrors of eugenic experimentation but have not eradicated ideas of race based biological difference in societies (i.e. the USA and Western Europe) that remain racialized (Saini, 2019). Modern examples of gynecological abuses and the use of race-based data correction are two contemporary instances where Black and Indigenous People of Color have been harmed by these lingering ideas (Montgomery, 2016; Aguilera, 2022; Vyas et al., 2020). The question then becomes how to educate current students of science to examine scientific findings for bias and how to train future scientists, medical professionals, and educators to avoid the incorporation of racially biased ideas into science and medical research methodologies, data interpretation, and clinical practice in professional settings.

The history of biological racism makes an awareness of the social impact and implications of a scientific work and the potential biases of its authors critical to appropriately understanding, interpreting, and using scientific findings responsibly (Nature, 2022). For science and medicine concerned with human health and behavior, societal and unconscious biases may reinforce or worsen existing inequities and discriminatory practices based on lingering myths about biological distinctions between different groups of people, e.g. races (Graves and Goodman, 2022; Dasgupta, 2020; Manali, 2018; Donovan et. al, 2019; Hind, 2023; Green et al., 2020). We posit that when scientists are considered first as people who possess their own backgrounds and biases, we may notice bias influencing their methodological choices and scientific conclusions, even when a work is technically sound. We created a teaching tool Finding inEquity in LIterature and eXperimentation (FELIX) to systematize the detection of bias and subjectivity in scientific communications.

### Theoretical Framework: Constructivist/Active Learning

The educational model used by our team was based on the Constructivist/Active Learning Theoretical Framework (Brandon & All, 2010). A constructivist approach uses instructional design where learning is framed as the development of personally meaningful understandings of content, developed through interactions with tools and others in a social context. Simply put, constructivism states that people construct their own understanding and knowledge of the world through experiencing things and reflecting on those experiences (Bereiter, 1994).

Honebein (1996) summarized seven pedagogical goals of constructivist learning environments as:

1. To provide experience with the knowledge construction process (students determine how they will learn).
2. To provide experience in and appreciation for multiple perspectives (evaluation of alternative solutions).
3. To embed learning in realistic contexts (authentic tasks).
4. To encourage ownership and a voice in the learning process (student centered learning).
5. To embed learning in social experience (collaboration).
6. To encourage the use of multiple modes of representation, (video, audio text, etc.)
7. To encourage awareness of the knowledge construction process (reflection, metacognition).

These pedagogical goals highlight a shift away from prior conceptions of teaching and learning where the teacher (i.e., expert) passively presents course materials to students who are empty vessels waiting to be filled. Rather, a constructivist approach acknowledges that learners are often confronting current understandings in light of new situations, leading to either a confirmation of what they know or the need to change or assimilate this new information. The modification of knowledge arises from the active process of applying current understandings, noting new elements in novel learning experiences, and adjusting based on the consistency of prior and emerging knowledge (Phillips, 1995).

### Theoretical Framework: Critical Reading

The importance of interpreting written texts in terms of authorship and societal influence has long been at the heart of critical pedagogy (Friere, 1985) and specifically critical reading pedagogies (Wolf and Barzillai, 2009). These frameworks are well established within education (Molden, 2007) and the social sciences (Van, Li and Wan, 2022; Jewett, 2007; Jensen and Scharff, 2019). The process of gathering information to place a text in its broader context to better understand it (sometimes called “deep”, “slow”, or “close” reading) requires time and effort. While several strategies have been proposed in the context of other disciplines (Jensen and Scharff, 2019), a formal or rigorous approach to performing this type of reading in the biological sciences is not widely practiced. In the sciences, critical reading is often presented in terms of scientific or information literacy with regard to public consumption of scientific texts (Priest, 2013). Students of biology and medicine learn methods for the technical reading and analysis of primary scientific literature with an emphasis on understanding the scientific context of an article, the methods used for the study, and the results of the study. However, it is rare to place these technical readings into a larger societal or historical context. The scientific method itself provides a framework that can be employed to systematize a deep reading of a text.

In scientific articles, important pieces of existing evidence are annotated with citations that provide the basis for the validity of a claim or assumption. However, while the same approach may be employed in fields and contexts outside scientific publications, this convention of explicitly citing prior evidence within a text is not as commonplace. It may be unclear in a text that a claim requiring substantiation has even been made, which statements are opinions vs facts, etc, thus making it difficult to know which statements to investigate. An intuition and sensibility for detecting aspects of a text that are biased or unsubstantiated may be developed with practice. FELIX combines these ideas of developing a systematic approach to critical analysis of text and using a standardized set of “measurements” into a tool for identifying and understanding a text with the goal of identifying potentially hidden biases. FELIX uses a three-phase progression: (i) Annotation, (ii) Analysis and (iii) Synthesis. Completion of each phase builds on the step before and guides the student from a detailed understanding of the text to a more conceptual understanding (Figure 1). In the annotation phase, students use a short vocabulary list of terms related to bias in scientific writing to highlight passages in the text. During the analysis, students answer critical analysis questions about the text and its authors to place the work in a social and ethical context. Lastly, via the Synthesis, students bring together elements from the annotation and analysis phases to form a unified argument about the text with a focus on its relationship to social or ethical context. We view this approach as key to placing modern research that deals with human health, genetics, and genomics, in both the context of current society and in relation to the history of biological racism and the subsequent harms that have been caused in the name of science.

**Figure 1:**
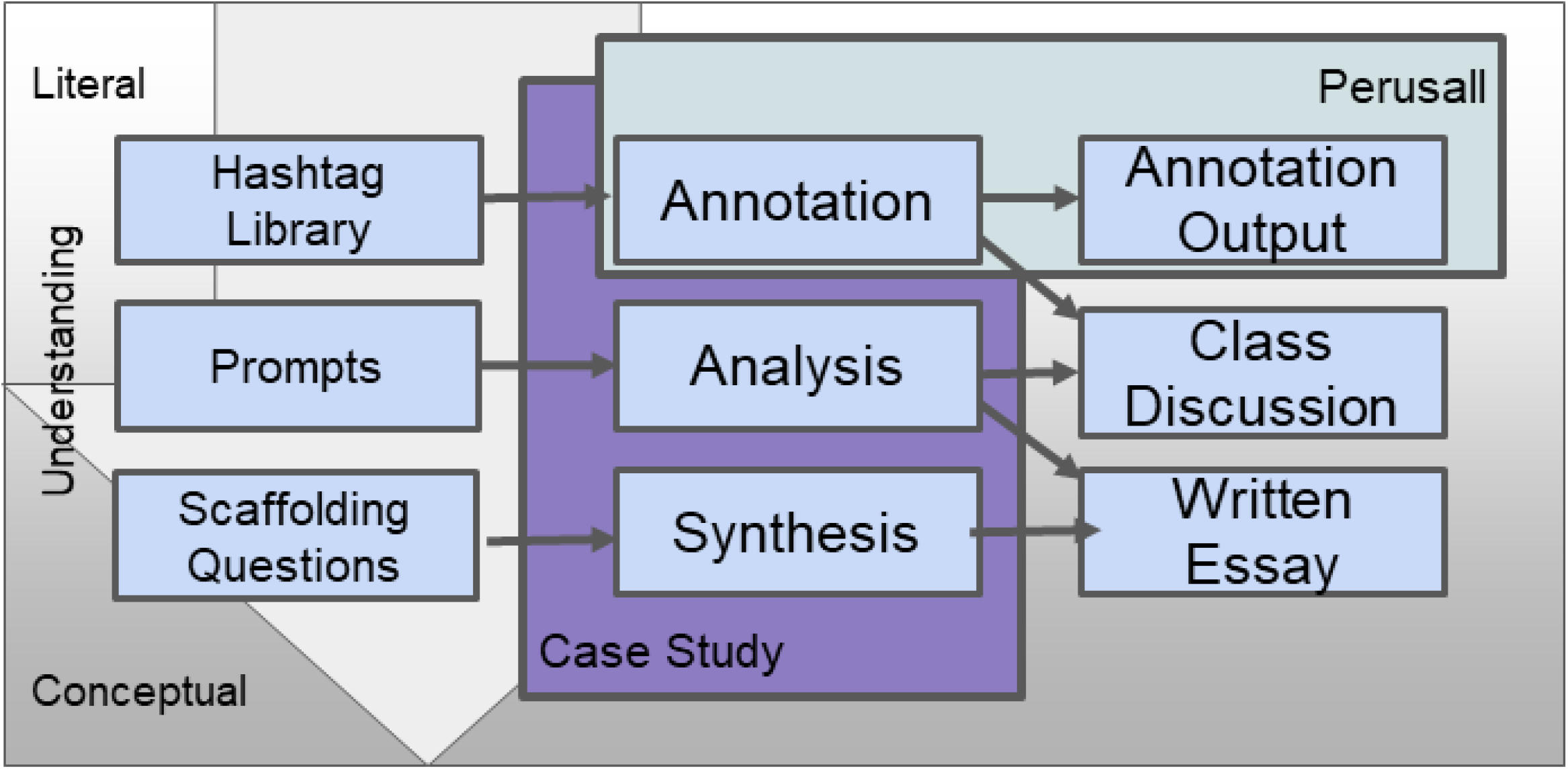
FELIX, a teaching tool for Finding inEquity in LIterature and eXperimentation. The conceptual framework and practical implementation of FELIX via the three phases – annotation, analysis, and synthesis.

### Research Questions

Here, we present the bias-detecting reading tool FELIX and results from its use in the context of an upper-level undergraduate biology course on institutional racism in health and science. Our instrument is intended to encourage students to deeply engage with course reading and was developed to achieve the following learning goals (i) develop student critical analysis skills when reading texts, (ii) help students build an intuition for identifying bias and opinions in texts, (iii) measure how student skills from (i) and (ii) change over the course of a semester, and (iv) create a dataset of annotations for a corpus of documents that capture the specific instances of bias-related aspects of the component texts. Our specific research questions revolve around evaluating item (iii), the measurement of changes in student skills over the course of semester. To specifically address this learning goal, we designed our approach with following research questions in mind:

Research Question 1: How do the perceptions and critical assessments of students change throughout the course of a semester using FELIX? We will address this question using quantitative analysis of the in-text annotations produced by four cohorts.

Research Questions 2: How do students enrolled in the Institutional Racism in Health and Science course at Boston University perceive the usefulness of FELIX? We will address this question using qualitative analysis of anonymous surveys from one cohort.

## METHODS

### Positionality

In relation to the proposed work, we are driven to dismantle and diversify the manifest, monolithic culture of whiteness in science. Although our white team members are beneficiaries of that culture, we recognize the toxic and counterproductive consequences of the exclusionary atmosphere it sustains. We are biased (we think positively) toward viewing systems through an antiracist lens, which may influence the design and interpretation of this work. As a team, we span a range of identities that has enabled us to reveal and mitigate many of our individual biases and blind spots. Each of us also sits at our own intersection of our various identities, providing intersectional perspectives on the issues we tackle. Over our years of work together, our team has built trust and cohesion that has been essential to supporting each other and fortifying this highly transdisciplinary, conceptually and emotionally challenging work on the legacy and reality of racism in our society. However, for all of our collective strengths and diversity, we recognize that many experiences and perspectives are not represented on our team. As faculty and staff at a major university in an affluent US city, we occupy a position of privilege which limits our ability to see the world from many perspectives. We further acknowledge that there are many perspectives which we have never encountered or imagined.

### Methodology

The researchers applied GTM (Grounded Theory Methods), an iterative qualitative approach designed to generate theory from data collected through interviews, observations, focused discussions, and document analysis. This choice was due to the complex and intertwined nature of the use of FELIX in a college course, the novel nature of our instrument, and the need to build theory based on empirical data collected during the semesters of use. From a constructivist approach, meaning making is a complex and iterative process and GTM aims to systematically collect and analyze data using an interpretive lens when the learning does not clearly include variables that can be statistically linked (Corbetta, 2003).

Researchers developed initial codes by parsing the data into segments, identifying key words, and concepts. Categories based on similarities and differences were developed from which themes and relationships were identified and then associated with categories, causal conditions, and consequences. Every new iteration of the course provided opportunities for theoretical sampling: researchers selected additional data to refine the emerging explanatory model (Charmaz & Bryant, 2019). Guiding their interpretation was the recognition that their own background and biases influenced their analyses. To counter potential pitfalls, the researchers constantly reflected on their own interpretations and biases, while raising questions and seeking alternative explanations. Their explicit goal was to center multiple perspectives, give voice to marginalized voices, challenge dominant narratives, and avoid imposing a single interpretation of the data (Charmaz & Bryant; Corbin & Strauss, 2008).

### Data Collection

#### Student Cohort and Survey Collection

Students enrolled in the course BI/BF510 Institutional Racism in Health and Science were surveyed anonymously as part of the university course evaluation process. The study cohort consisted of students of junior and senior undergraduate standing as well as graduate MS or PhD candidate status from five semesters of the course for qualitative data (Total N=78; F21 (N=18); S22 (N=16); F22 (N=12); S23 (N=22); F23 (N=10) and one semester of the course for quantitative data (F23, N=13). The student pool is enrolled in degree granting programs in Biology and Bioinformatics. Students were given 5 minutes at the beginning of class on the first and final days of the course. Surveys was generated using Qualtrics with QR link generated via open web tools (qr.io); thereby adding accessibility via mobile devices. Students used the projected link or QR code to complete the assessment. Pre and post survey questions included perceptions of science and racial bias. The post survey also asked about student experiences with FELIX. The assessment used a simple 1-5 scale indicating level of agreement (disagree completely – 1; disagree somewhat – 2; not sure/don’t know -3; agree somewhat – 4 and agree completely – 5). A condition of the assessment is that instructors do not assess the exit results until after grades are submitted for the semester.

#### Application of FELIX to Course Readings: The Hashtagulary and Annotation Analysis

FELIX is composed of three phases: 1) annotation, 2) analysis, and 3) synthesis. During the annotation phase, students use the Perusall web application (Clarke, 2019) to perform and record hashtag annotations (e.g. #opinion) of assigned articles throughout the semester. Students may use any hashtag they deem appropriate, but are instructed to first consider a standardized set of hashtags (the “hashtagulary”, portmanteau of hashtag and vocabulary) that we developed to label elements that might be related to bias. By annotating a text with a controlled vocabulary of hashtags, we can create a consistent dataset that enables meaningful quantitative analysis. The purpose of the hashtagulary is to enable meaningful textual analysis of annotations made by many people of the same text, thus enabling algorithmic characterization of the presence and specific passages that might contain bias. The hashtagulary is listed in Table 1.

**Table 1:**
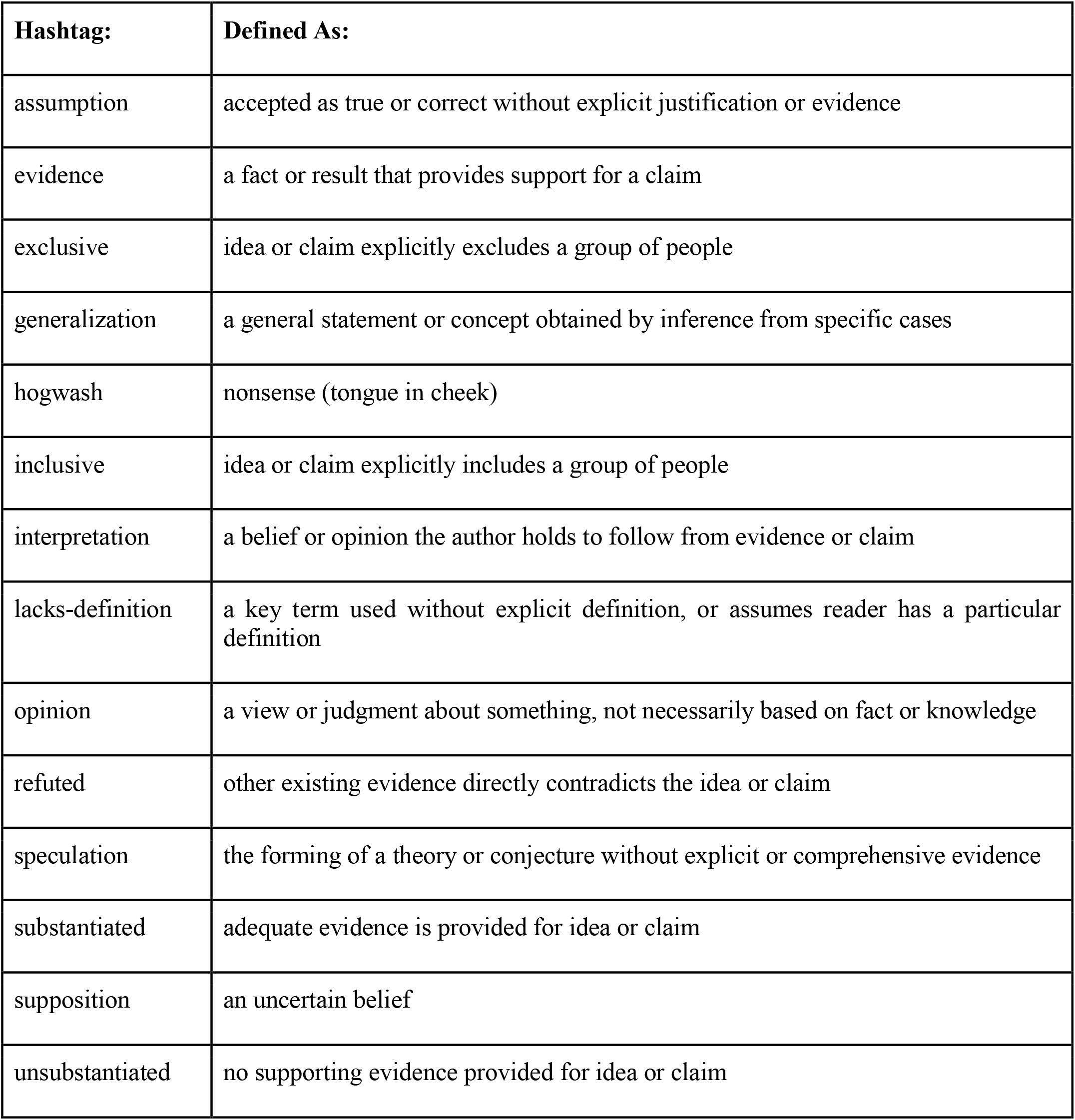
The Hashtagulary - a hashtag vocabulary defined as set of hashtags used by students to annotate texts. Students may use other hashtags as they see fit but are instructed to consider these definitions first.

Perusall allows export of all annotation data that includes date, article, annotator, annotation comment, and annotation position in the text. This data is then downloaded, cleaned, and analyzed to generate a processed dataset of various hashtag counts and locations. The cleaning and analysis are performed in Python and the current analysis is performed using Jupyter notebooks.

#### Data Sharing

Jupyter notebooks were used for hashtag analysis and can be found at the following locations: https://colab.research.google.com/drive/1aUl-mRKmqKl6ItIA9a3UyeqmuQl_5kZ3?usp=sharing

#### Quantitative Assessment of FELIX – Survey Analysis

Quantitative surveys were used to investigate the effect of the course on the effectiveness of FELIX as a teaching tool. Surveys were generated using Qualtrics with QR link generated via open web tools (qr.io); thereby adding accessibility via mobile devices. Students used the projected link or QR code to complete the assessment. Students were given 5 minutes at the beginning of class on the first day of the course for the entry survey and were sent the same survey after final grades were submitted via email. Pre and post survey questions included perceptions of science and racial bias. The post survey asked about student experiences with FELIX. The assessment used a simple 1-5 scale indicating level of agreement (disagree completely – 1; disagree somewhat – 2; not sure/don’t know -3; agree somewhat – 4 and agree completely – 5). A condition of the assessment was that instructors do not assess the exit results until after grades are submitted for the semester.

#### Qualitative Analysis of Student Feedback – Student Questionnaire

Qualitative feedback from the students was collected from students each semester through an anonymous exit survey completed when the students handed in their final projects during finals week. A condition of soliciting feedback was that instructors do not assess the exit results until after grades are submitted for the semester. We posed the following question to the cohort regarding the use of FELIX in the course - “Did you find the instrument to be effective at understanding bias in a text?” Students were asked to further elaborate on why or why not they found the instrument to be helpful. Students were also asked to specify which phase of FELIX they found to be the most useful to their learning. We compared terms in the evaluations that indicated both positive student experiences and conversely, student confusion. Criterion for the different themes were as follows - positive themes included language that indicated a deeper understanding of the material; improved reading comprehension; new perspectives, increased critical thinking, and intention to use skills from the instrument beyond this classroom; negative themes – included expression of confusion and dislike of any aspects of the process; neutral themes – were the absence of positive or negative themes.

## RESULTS

### Hashtag and Annotation Analysis

The data from the annotation phase of this study was used to construct heatmaps showing key moments in the text. A heatmap is a data matrix where coloring is used to illustrate an overview of the numeric differences. Our heatmaps were created in Python to note which hashtag students used to annotate the text (on the vertical axis) and the location of that annotation in the overall journal article (on the horizontal axis). Figure 2 shows student annotations for the article Race Crossing in Jamaica [13]. The top left (A) provides overall counts of the annotations used, the top right shows (B) shows how a correlation matrix of the hashtags used, and the bottom heatmap (C) shows specific locations in the text where annotations were used. These visualizations highlight the effectiveness of our implementation of FELIX in this course as they provide a window into what students choose to focus on and how they are interpreting that passage of text. A heatmap was created for each article, allowing our teaching team to assess how students were grasping the course content and inform the focus of our in-class discussions.

**Figure 2:**
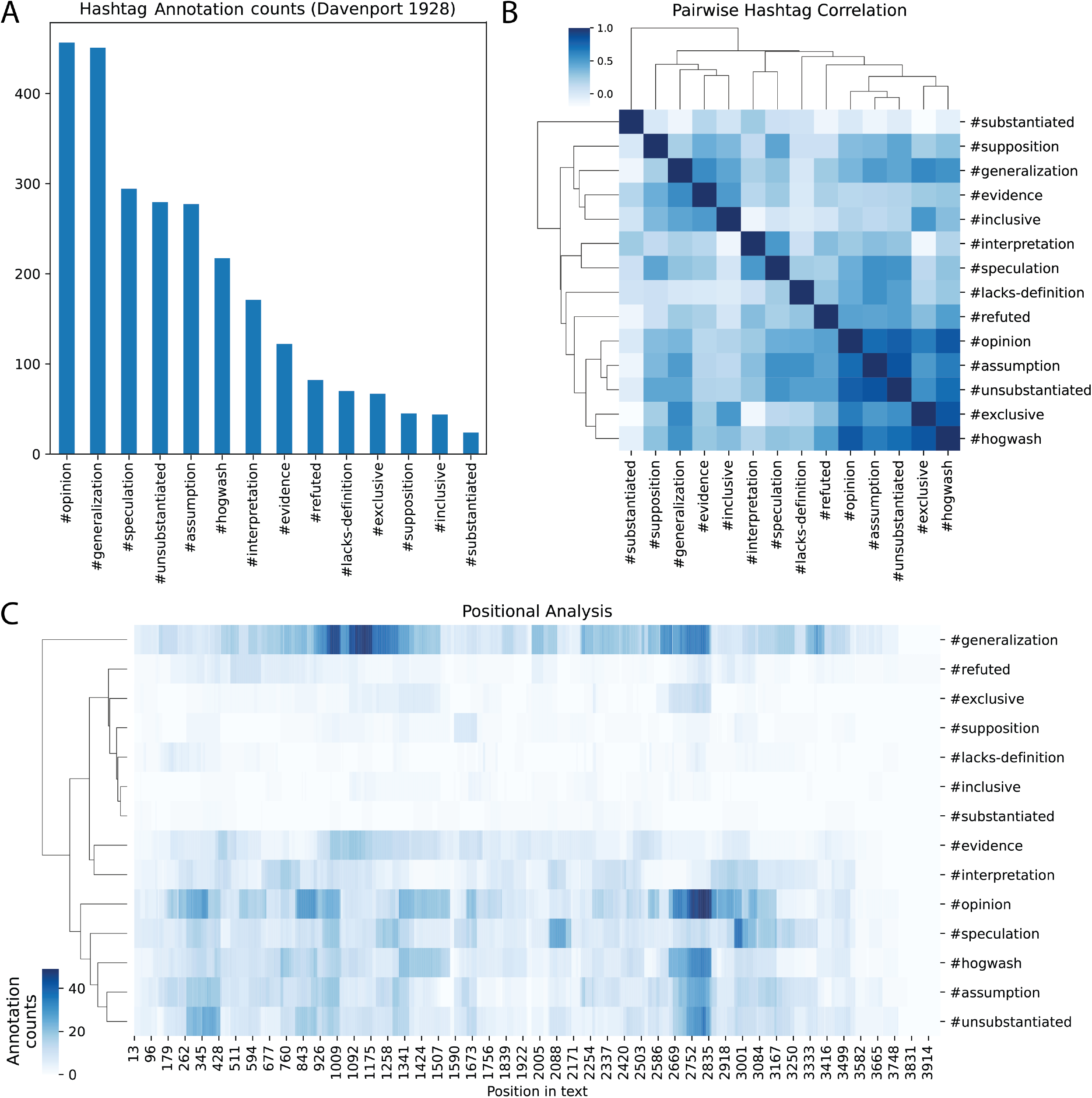
Annotation analysis of Davenport “Race Crossing in Jamaica” 1928 from four semesters. A) Total count of hashtag annotations. B) Correlation of pairwise hashtags, i.e. hashtags frequently annotated to the same passage have high correlation. C) Positional analysis showing passages in text annotated with corresponding hashtag. Left to right is position in text, darker colors indicate more annotations of a hashtag to that location. Text passages (1,2,3) with notable annotations (#generalization, #opinion, #speculation, respectively) are quoted from Davenport as indicated.

We also examined the annotation for evidence of course-wide trends in student learning and development. After cleaning, students used 541 unique hashtags (14 are in the hashtagulary) across 13,231 annotations made by 82 students. 68% of the annotations made utilized the hashtagulary hashtags. We compared the annotation count for hashtagulary or student created hashtags and found some hashtags were more frequently annotated than others, where some student created hashtags were used with greater frequency than the standard set (Figure 3A). When hashtags patterns were compared with the chronological course as measured by number of days into the semester (to normalize across all four semesters) students consistently used hashtagulary annotations throughout but used increasingly more of their own hashtags as the semester progressed (Figure 3B). We interpret this to mean students gained more comfort and skill using FELIX annotations with practice. Finally, we performed a sentiment analysis of annotations by manually annotating each hashtag with either a -1, 0, or 1 based on whether the hashtag expressed negative, neutral, or positive sentiment, respectively. For example, hashtags #injustice, #opinion, and #substantiated were labeled as -1, 0, and 1 respectively. We see a trend toward positive sentiment across the semester, which we interpret as due to the course readings being published closer to the present as the semester proceeds.

**Figure 3:**
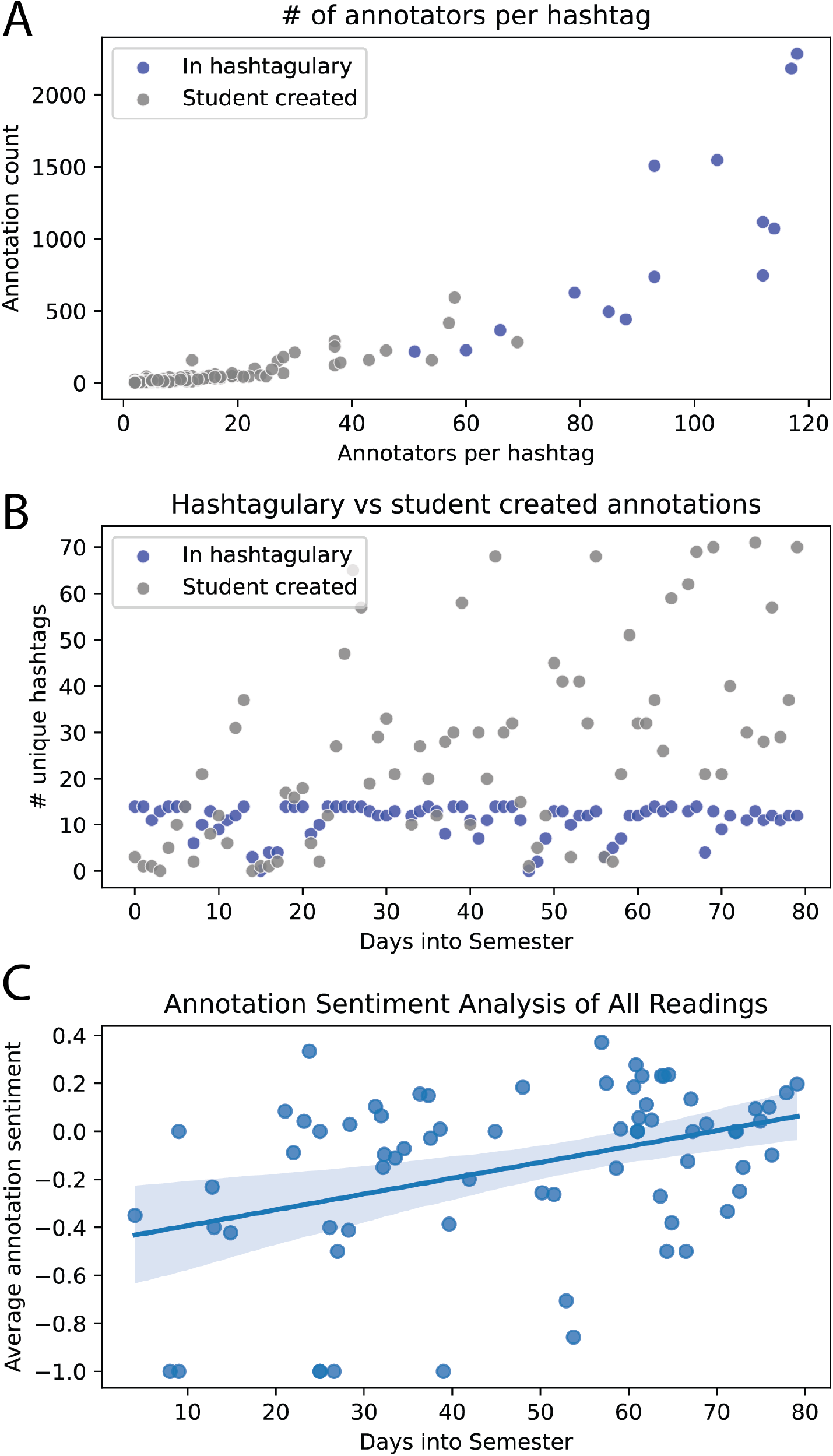
Annotation Trends. A) Number of annotations vs number of students using each hashtag. B) Number of unique hashtags used in annotations by all students vs chronological day in the semester. C) Mean sentiment for all hashtags in all students for each reading across each semester.

### Student Perspectives and Analysis of FELIX

Quantitative feedback from the students in IRHS indicated an overall positive viewpoint on FELIX as a reading method (Table 2), with students agreeing statements positing the usefulness of each phase of FELIX (Annotation; Analysis; Synthesis). The effectiveness of FELIX as providing a new perspective to understanding the intersection of race, genetics, and biology was also indicated in the results (Figure 4). Student perspectives shifted over the course of the semester with less agreement at the end of the semester with statements linking race with genetics and biology (orange dots, Figure 3). Furthermore, student perspectives about the objectivity of science and scientists indicated a greater awareness of the bias present in both the system and the people in the system (orange dots, Figure 3).

**Table 2:** Results from Quantitative Evaluation of FELIX Post Semester (F23; N=8). The assessment used a simple 1-5 scale indicating level of agreement (disagree completely – 1; disagree somewhat – 2; not sure/don’t know -3; agree somewhat – 4 and agree completely – 5). Higher scoring indicates MORE agreement. (N=8).

| **<u>Statement</u>** | **<u>Mean</u>**<br>**<u>(5=agree</u>**<br>**<u>completely)</u>** | **<u>Standard</u>**<br>**<u>error</u>** |
| --- | --- | --- |
| I found completing annotations to be useful to understanding the assigned text. | 3.9 | 0.41 |
| I found completing the analysis questions in groups during class discussion time to be useful to understanding the assigned text. | 4.7 | 0.15 |
| I found completing the analysis questions in the case studies to be useful to understanding the assigned text. | 4.7 | 0.15 |
| I found the synthesis step (case study) useful to understanding the assigned text. | 4.3 | 0.34 |
| I found the reading approach (FELIX) to be useful/helpful. | 4.6 | 0.24 |
| Using the reading approach (FELIX) in class put science papers into larger context. | 4.7 | 0.23 |
| The authorship of a paper can influence its conclusions. | 5.0 | 0.0 |

**Figure 4:**
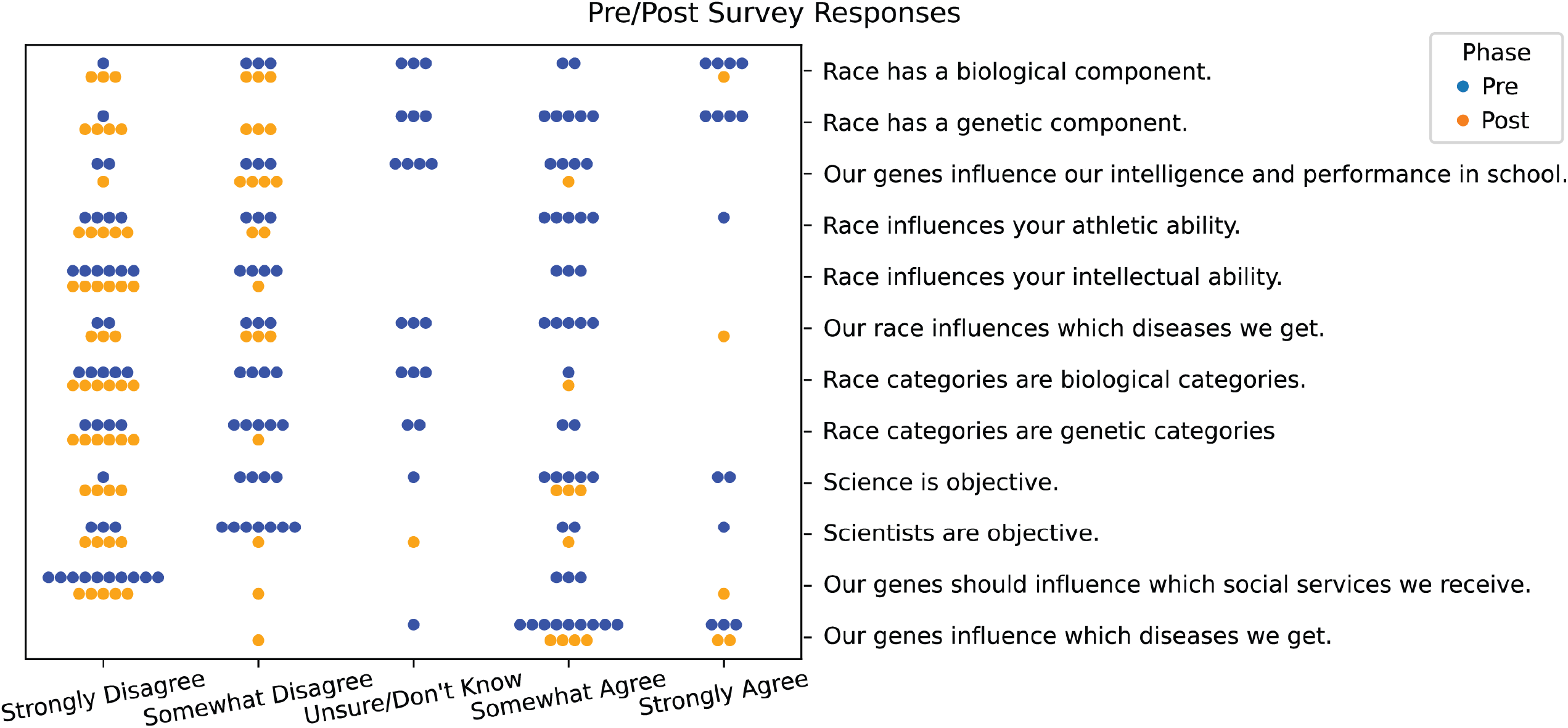
Students’ perceptions of key course concepts at the beginning of the semester (Pre; blue dots; N=15) and at the end of the semester (Post; Orange dots; N=8).

We collected additional qualitative feedback from the students in IRHS indicating the effectiveness of FELIX in improving student learning and in providing a new perspective to training in the biological sciences (Table 2). We posed the following question to the cohort after submission of the final project at the end of the semester - “Did you find the instrument to be effective at understanding bias in a text?” Most students from across all four cohorts answered “yes” to the effectiveness of the instrument (87%, 68/78 responses), with 12% of the cohort responding neutrally (9/78) and 1% responding negatively (1/78).

Students were asked to further elaborate on why or why not they found the instrument to be helpful. For example, one student responded:

> The instrument was incredibly effective at understanding bias in a text. I felt that using the three steps of the instrument made you engage with the readings in a way that just reading it would not. I particularly thought that the annotations were the most impactful in making me understand bias in the readings. (S8; 12/13/21)

This student highlights how FELIX allowed them to engage with the readings in a way that allowed for a deeper understanding of the course materials. Another student shared how the steps of FELIX helped them in this course and how they plan to apply these in their future:

> I absolutely will be using the techniques I learned while reading papers in the future; it made me look at the papers more deeply than just absorbing the scientific information presented. I felt the annotation and analysis were very helpful and the analysis questions are something I can use in my everyday life while reading papers to think about who is included/excluded. (S5; 12/18/2022)

We compared terms in the evaluations that indicated both positive student experiences and conversely, student confusion. Positive themes we saw indicated a deeper understanding of the material (N=20); improved reading comprehension (N=6); new perspectives to learning the material (N=6), increased critical thinking (N=6) and intention to use skills from the instrument beyond this classroom (N=4) (Table 2). Because these labels were not overlapping, this indicated that 42/68 evaluations had language specifically denoting new skills attained by the students from using this tool.

Students were also asked to specify which phase of FELIX they found to be the most useful to their learning. Students specifically cited more utility for annotation (N=32) vs Analysis (N=30) vs Synthesis (N=20). These data are more reflective of themes that developed in the analysis. First, students indicated a positive appreciation of the application of the phasewise process of FELIX (N=7), in which the annotation and analysis phases were linked together. Secondly, those students indicating appreciation of analysis also indicated appreciation for digging deeper into the backgrounds of the authors of scientific papers (N=3).

## DISCUSSION

This study contributes a unique pedagogical approach to reading scientific articles in the context of upper level, undergraduate biology courses - one in which the history of biological racism is brought to bear in looking for lingering racial bias in genetics, genomics, and human health literature. Our approach is autodidactic in nature, with student learning that is generated from the engagement of the students with the reading process via the FELIX framework. Additionally, knowledge is generated about the texts at hand via the annotation process that is carried out during the annotation step of the instrument and generates a dataset of student hashtags for each reading throughout the semester (Figure 3).

Our quantitative analyses of student perceptions from one cohort (Fall 2023) suggests the effectiveness of FELIX in changing student viewpoints on the objectivity of scientists and the link between biology and racism over the course of one semester. However, there are several limitations to the initial analyses. Since the quantitative survey was a pilot (N = 13 pre / 8 post), the sample size for the data is small and requires larger numbers to corroborate the pilot findings. Secondly, the pilot questions require revision to include control questions regarding student perception of learning. Currently, all the survey questions are stated positively and in a specific order. We would seek to include questions for which a negative or neutral response would be expected and to randomize the order in which questions are posed. The authors recognize the irony of trying to quantify student perceptions of their critical (and subjective) analyses of texts. It is inherently difficult to introduce a tool for critical thinking and to widen perspectives and then to assess that tool in an objective way. We are excited to refine our methods and expand our data collection in current and future cohorts to face these challenges.

Overall, qualitative student feedback also indicated FELIX provided a systematic approach that students found to be a useful tool for their learning. Students reported that it helped them to improve their reading comprehension, critical thinking skills, and in understanding the social and ethical implications of the research they were reading. Nonetheless, there are areas of our methodology that can be improved. Students indicated a preference for the earlier steps of the instrument - annotation and analysis. Feedback suggested that the current implementation of the synthesis phase could be improved to tie in the previous two phases more strongly and to engage students more effectively. When it came to the annotation process, students also indicated some confusion with the hashtags and feeling restricted by the vocabulary. Future iterations will work to make the hashtag vocabulary a more dynamic process with student input. Given the positive qualitative feedback collected over all five of the course offerings (N=78), we are currently working to develop shorter workshop versions of FELIX that can be offered on a condensed time frame compared to a full semester course.

Lastly, our assessment surveys - both qualitative and quantitative - have focused on perceived learning on the part of the students. While this may be a good way to indicate student appreciation for the material being learned; a more concrete assessment method would be ideal for determining if critical reading skills are indeed increasing from using FELIX. Future studies in which students do a pre and post annotation of articles is in the process of being designed and implemented to better quantify changes in use of hashtags and indication of biases before and after becoming familiar with use of our instrument.

The result of our pilot leads us to a lingering research question: does learning about the history of harms perpetrated by science and medicine due to racial bias change how scientists and medical professionals think? Does it generate more empathetic and aware practitioners who will carry out their work and research in ways that ultimately cause less harm? How can one quantify life-long effects from encouraging students to think about the social and ethical implications of their scientific research? Regardless of how we answer these questions, history shows us that bias can influence the design, conduct, and interpretation of scientific research. This history is a reason why learning how to become critical readers of scientific literature should be a key component of university teaching and learning. Future work will include building on this foundation to compare learning outcomes to cohorts with and without training using FELIX and tracking student use of FELIX in contexts outside of our course. Our piloting of this method and the preliminary indications of its efficacy lead us to put forward this approach as a universal method for training students in the reading of scientific literature and communications.

## ACKNOWLEDGMENTS

Thank you to the Lab of Daniel Segre at Boston University for discussion and helpful feedback on FELIX and the course, Institutional Racism in Health and Science, as well as for reading and discussing drafts of this manuscript. Thank you to all the IRHS510 students, who inspire and motivate us in our work every day.

