## Supplemental Table 1 for "Reading Outside the Lines: A Systematic Approach for Detecting Bias in Scientific Communications"

**Supplementary Table 1: Quantitative and Qualitative Survey Questions Used for the Pilot Evaluation of FELIX and suggested question revisions for future iterations.**

| | |
|---|---|
| | **QUANTITATIVE ASSESSMENT** |
| ***Section 1 (Pre&Post) – Questions Regarding Student Perceptions of Science and Medicine*** | **Instructions - How much do you agree with the following statements? (disagree completely – 1; disagree somewhat – 2; not sure/don't know - 3; agree somewhat – 4 and agree completely – 5).** |
| | Race has a biological component. |
| | Race has a genetic component. |
| | Our genes influence our intelligence and performance in school. |
| | Race influences your athletic ability. |
| | Race influences your intellectual ability. |
| | Our genes influence which diseases we get. |
| | Our race influences which diseases we get. |
| | Race categories are biological categories. |
| | Race categories are genetic categories |
| | Science is objective. |
| | Scientists are objective. |
| | Our genes should influence which social services we receive. |
| ***Section 2 (Post only) Questions regarding the use of FELIX three step approach for class reading.*** | **Instructions - Please indicate your level of agreement with each statement.** |

| | |
|---|---|
| | I found completing annotations to be useful to understanding the assigned text. |
| | I found completing the analysis questions in groups during class discussion time to be useful to understanding the assigned text. |
| | I found completing the analysis questions in the case studies to be useful to understanding the assigned text. |
| | I found the synthesis step (case study) useful to understanding the assigned text. |
| | Overall, I found the reading approach (FELIX) to be useful/helpful. |
| | Using the reading approach (FELIX) in class put science papers into larger context. |
| | The authorship of a paper can influence its conclusions. |
| | **Qualitative exit survey questions (long answer)** |
| Question 1: | Did you find the instrument to be effective at understanding bias in a text? Why or why not? Was there a specific step that was more useful than others (annotation vs analysis vs synthesis)? |
| Question 2: | Did you find the process of using the instrument to be valuable to preparing your final project? Why or why not? |
| | **Proposed Revisions of Qualitative Survey Questions** |
| | How would you describe your experiences as a student using FELIX to analyze text? What, if anything, did you find beneficial? |
| | What are your thoughts and feelings about the themes that emerged in your analysis? |
| | In your opinion, what are the biggest challenges facing students who plan to pursue careers in science or health? What are the biggest opportunities facing students who plan to pursue careers in science or health? |

| | |
|---|---|
| | How, if at all, did FELIX helped you to think about structural inequities in your aspiring profession? What, if any, questions does using this instrument raise for you about your ethical responsibility in your chosen profession? |
| | What, if anything, in the instrument be adjusted to better support your developing understanding of existing structural disparities in health and science? |
| | What, if any, other insights could you offer about the use of this instrument as a tool of analysis? |
